# Cardiac myocyte microtissue aggregates broadcast local field potentials

**DOI:** 10.1101/376418

**Authors:** Mijail D. Serruya, Suradip Das, Kritika S. Katiyar, Laura A. Struzyna, Justin C. Burrell, D. Kacy Cullen

**Author notes:** Co-First Authors (Mijail D. Serruya, Suradip Das). **Correspondence**: Dr. Mijail Serruya.

## Abstract

Muscle tissue has been exploited as a living biopotential amplifier to facilitate transduction of peripheral nerve signals into prosthetic control in patients with limb amputation. Here we sought to address the question of whether microscopically small volumes of muscle tissue could effectively broadcast field potentials to electrodes not immediately in contact with that tissue. Cardiac myocytes were grown as three-dimensional aggregates containing 10^5^ cells comprising a volume of approximately 0.065 mm^3^ (~500 μm in diameter) atop multi-electrode arrays. In addition to the expected spontaneous contraction potentials detected using electrodes in direct contact with the myocytes, potentials could also be detected on distant electrodes not contacting the aggregates. Specifically, while both dissociated and aggregated cardiac myocyte cultures generated spontaneous contractions that could easily be recorded from underlying multi-electrode arrays, only aggregated myocyte cultures generated signals detectable several millimeters away by the electrode grid floating in media. This confirmed the ability of micro-volumes of aggregated muscle tissue to broadcast readily detectible signals. The amplitude of the potentials generated by the aggregates decreased exponentially with distance. The aggregates were sensitive to pharmacologic modification with isoproterenol increasing contraction rate. Simultaneous recordings with electrodes in physical contact to the aggregate and with electrodes several millimeters away revealed that the aggregates function as amplifiers and low-pass filters. This study lays the groundwork for forging myocyte aggregates as “living amplifiers” for long-term neural recording in brain-computer interfaces to treat neurological disease and injury.

## 1 Introduction

Animals have evolved muscle tissue both to achieve locomotion as well as a wide range of internal functions including circulation and digestion. In the process of achieving mechanical action, muscle tissue also effectively acts as a living biopotential amplifier, such that generated myopotentials are several orders of magnitude greater in amplitude than the neural potentials emerging from the input nerves. Bioengineers have exploited the amplification properties of muscle tissue to facilitate recording of signals from the brachial plexus and peripheral nerves in patients with amputation. For instance, in targeted muscle reinervation, remaining nerve branches of the brachial plexus are repositioned over the pectoralis major in an upper extremity amputee, such that the peripheral nerves spontaneously innervate thin strips of the pectoralis. Electrodes placed on the skin surface of the chest in turn detect the brachial plexus branch-triggered muscle action potentials in the pectoralis and these signals can be decoded into intended forearm movements to control prosthetic limbs with sufficient dexterity to perform activities of daily living [1]. An alternative approach has been to biopsy small flaps of muscle tissue and fold them over the cut ends of the peripheral nerves (like a “taco”). By providing the traumatized nerves a new target, this technique has been shown to prevent the development of neuromas and hence spare patients considerable discomfort [2]. In regenerative peripheral nerve interfaces, electrodes are placed in proximity to these miniature muscle grafts to pick up these myopotentials and similarly decode them into forearm instructions to control a prosthetic [3]. By “plugging in” nerves to endogenous or augmented muscle tissue, the surgeons have dramatically simplified the technical recording challenges that bioengineers must overcome to provide a reliable medical device.

These promising innovations raise the question: how small can we go in terms of using muscle tissue to amplify neural signals? Within the human body, the 1-mm long stapedius is the smallest skeletal muscle, and smooth muscle tissue dwindles in volumes of sub-cubic-millimeter scale; other animal species have muscular units operating at micron scales. As the volume of muscle tissue decreases, however, the amplification properties also decline, and to our knowledge the ability of muscle tissue to broadcast signals at the micron scale has never been systematically explored. Following the precedent set by macroscopic peripheral nerve-muscle amplification for prosthetics, we anticipate that micro-scale equivalents may likewise use motor neurons coupled to skeletal myocytes.

While functional neuromuscular junctions spontaneously form in motor neuron-skeletal myocyte co-cultures [4], [5], the ability to ensure that a sufficient number of such junctions form to reliably induce a large myopotential has not been determined. To render the design challenge tractable, as a proof-of-concept we chose to begin with myocyte-only constructs. Since skeletal myocytes tend to be quiescent, and since we wanted to avoid confounds and complexity that would be introduced by optogenetics, we decided to culture cardiac myocytes because they spontaneous contract, and via gap junctions, coordinate this contraction across cells. Therefore, the following study used tissue culture and electrophysiological techniques to characterize the electrophysiological properties of novel three-dimensional (3D) microtissue comprised of engineered cardiac myocyte cell aggregates.

## 2 Methods

### 2.1 Rat cardiac myocyte harvest and forced aggregate culture

All procedures involving animals were approved by the Institutional Animal Care and Use Committee of the University of Pennsylvania and followed the National Institutes of Health Guide for the Care and Use of Laboratory Animals [6]. Hearts were harvested from E16 Sprague Dawley rat pups. All harvest procedures prior to dissociation were conducted on ice. The pups were decapitated, and a small incision was made to expose the thoracic cavity. The heart was freed from surrounding tissue, taken out and kept in HBSS on ice. An average four hearts were taken for subsequent primary cultures. The hearts were digested in 0.05% Trypsin-EDTA for 10-15mins at 37°C. The tissue digestion was terminated by addition of HBSS+10% FBS. The tissue was then triturated thoroughly to form a homogenous solution. 4% BSA was added to the bottom of the tissue solution as a cushion before being centrifuged at 300g for 10mins. The cell pellet was dissolved in cardiac media (78% DMEM-High glucose + 17%Medium-199 + 4% Normal Horse Serum + 1% Penn-Strep) to obtain 7x10^6^ cell/ml. To create aggregates, dissociated cardiac myocytes were plated in “pyramid” shaped polydimethyl siloxane (PDMS) wells using techniques similar to those described previously [7]–[10] Cell suspension volume of 14 μL (~10^5^ cells) was added to each pyramid, and centrifuged at 1500 RPM for 5 min. The wells were then flooded with cardiac media and incubated for 24 hours to allow the aggregates to form. A total of 18 cultures were examined for this study: n=13 using engineered myocyte aggregates and n=5 using planar myocytes, broken down as detailed below.

### 2.2 Planar multi-electrode arrays

Multi-electrode recordings were made with the MED64 system (AutoMate Scientific, Inc, Berkeley, California). The MED probe contains 64 electrodes, each 50 μm across, in an 8×8 grid with interelectrode spacing of 75 μm, and an average impedance of 10 kΩ for a 1 kHz 50mV sinusoid.

### 2.3 Floating multi-electrode grids

A planar 30-channel mouse EEG (H32, serial number D70C, NeuroNexus Technologies, Inc, Ann Arbor, Michigan) was used to record local fields at a distance from the cultured cells. The mouse EEG electrodes are platinum embedded in a polyimide substrate with an array thickness of 20 μm; the impedance ranges from 5 to 20 kΩ for a 1 kHz 50mV sinusoid. The mouse EEG is mounted on a micro-manipulator and lowered to float on the surface of the media (Figure 1).

**Figure 1.**
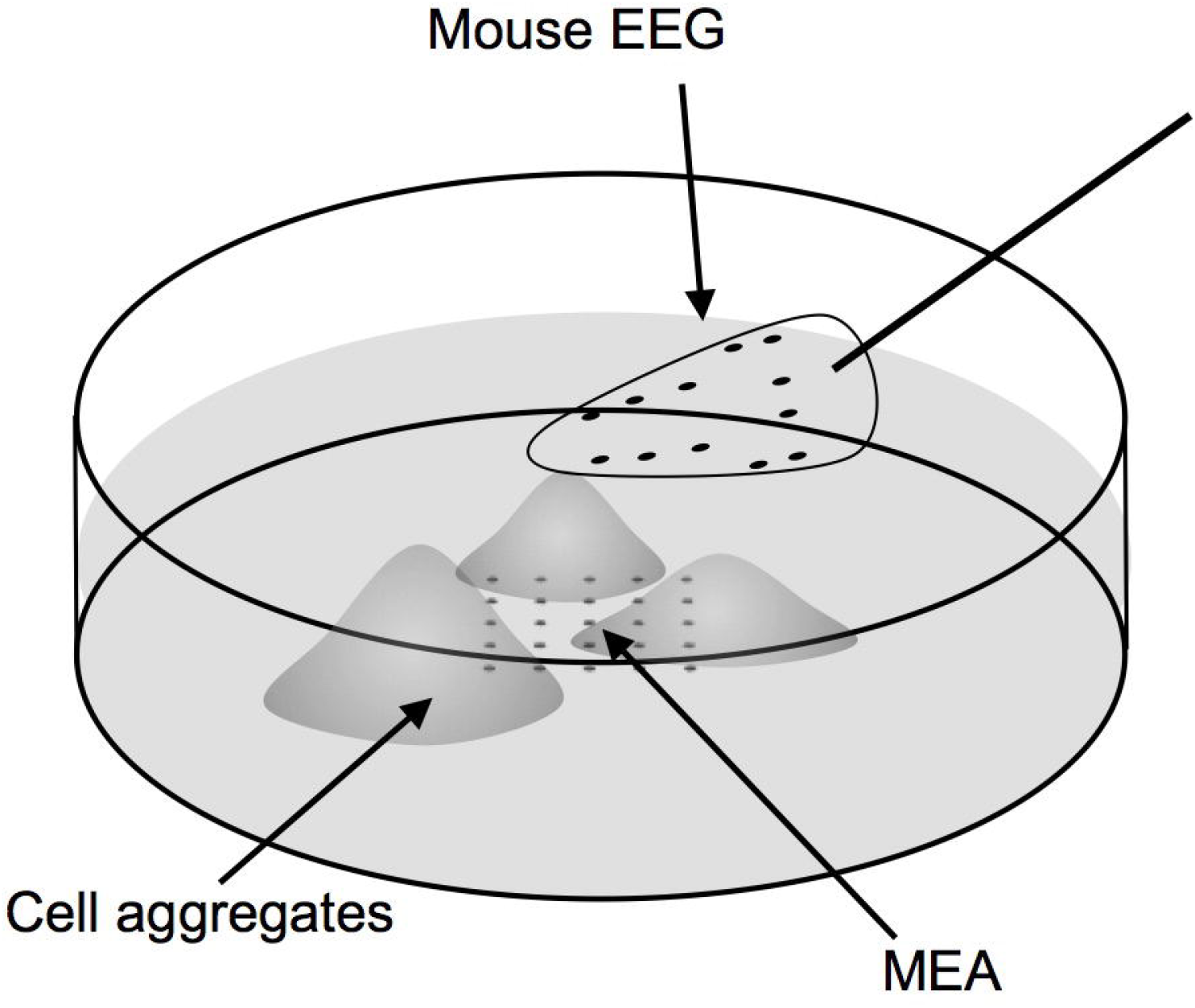
The experimental setup includes cardiac myocytes grown as forced aggregates atop a rigid MED64 high-density multi-electrode array (MEA). A flexible “mouse EEG” is floated in the media above the cultured myocytes. Each electrode array is in turn connected to a 32-channel digital head stage that sends signal into the amplifiers and recording computer.

### 2.4 Cardiac myocyte culture on multi-electrode array dishes

The MEA dishes were coated overnight with 20μg/ml of poly-ethyleneimine followed by 2 hours of treatment with laminin (20μg/ml). For dissociated cultures, 150μl of cell solution (7x10^6^ cells/ml) was directly added on the MEA dish. The plate was filled up with 2 ml of cardiac media. The cardiac myocyte aggregates (3-4) were pipetted out of the pyramid wells and carefully placed on the electrode array region within the MEA dish. The aggregates were allowed to adhere for 3-4 hours, filled up with 2ml of media before moving the MEA into a cell culture incubator. Both the dissociated as well as aggregated cardiac myocyte cultures on the MEA were grown for 5 days *in vitro* (DIV) with regular change of media before performing electrophysiological recordings. Sister cultures were maintained in identical conditions and grown on PDL-Laminin coated petri dishes. The cardiac myocytes were treated with 10μM of isoproterenol (isoprenaline hydrochloride, a β-adrenoreceptor agonist) in cardiac media to induce an *in vitro* correlate to tachycardia.

### 2.5 Recording of electrical activity

The MEA dishes containing dissociated or aggregated cardiac myocyte cultures were placed in an adapter frame (item AL-MED-C03, AutoMate Scientific) to which one or two 32-channel digital headstages were attached in turn connected to cables delivering data into the neural data acquisition system (Digital Lynx; Neuralynx, USA). An alligator clip was attached to the wall of the dish such that it was submerged in the media: the other end was attached to the reference connector of the adapter frame. For recording potentials in saline from a distance, the mouse EEG was floated in the media above the cultured cardiac myocytes and plugged into a 32-channel digital head-stage, also linked to the neural data acquisition system; this head-stage also had a reference wire that was tied to a second alligator clip also placed in the media. The data acquisition system acquires samples at 30 kHz. Data was analyzed in Matlab (The Mathworks Inc., Natick, Massachusetts, USA). Electrical activity recording studies were performed on both planar (n=1 for EEG only and n=1 for MEA+EEG) and aggregate (n=4 for MEA only, n=3 for EEG only, and n=3 for MEA+EEG) cardiac myocyte cultures.

### 2.6 Immunocytochemistry and imaging of cardiac tissue cultures

All cardiac cell/tissue cultures were routinely imaged using phase contrast microscopy techniques on a Nikon Eclipse Ti inverted microscope with Nikon Elements Basic Research software. For immunocytochemistry analysis, 5 DIV cultures were fixed in 4% formaldehyde for 30 min, rinsed in phosphate buffered saline (PBS), and permeabilized using 0.3% Triton X100 plus 4% horse serum for 60 min. Primary antibodies used to identify cardiac myocytes were added (in PBS + 4% serum solution) at 4°C for 12 hrs. Rabbit anti- cardiac troponin T (Abcam ab45932) was used to identify a cardiac specific tropomyosin binding subunit of troponin. After rinsing, Secondaries (1:500 in PBS + 4% NHS) were applied at room temperature for 2 hours (Alexa 488 donkey anti-rabbit IgG). Dissociated and aggregated cultures (n=3 each) were fluorescently imaged using a laser scanning confocal microscope (LSM 710 on an Axio Observer Z1; Zeiss, Oberkochen, Germany). For each culture, multiple confocal z-stacks were digitally captured and analyzed.

## 3 Results

Primary cardiac myocytes grown in novel 3D spherical aggregates were compared to those grown in traditional planar cultures based on growth, phenotype, and functional characteristics. Specifically, dissociated cardiac myocytes were subjected to “forced aggregation” using gentle centrifugation in custom-built pyramidal micro-wells, and the resulting engineered aggregates were comprised of 10^5^ cells ranging in diameter from 500 – 800 μm. Given a diameter of 500 μm, and assuming a roughly spherical geometry at the time of plating, the initial volume of these aggregates was estimated to be 0.065 mm^3^ (0.065 μL or 6.5x10^-5^ mL).

The cardiac myocytes in both aggregate and planar culture were observed to beat spontaneously from day 1 in culture. By 5 DIV dissociated planar cultures exhibited sporadic beating of individual cells whereas the aggregates were observed to beat synchronously as a single unit (see Supplementary movies). The cells under both conditions exhibited a healthy morphology and had robust expression of cardiac troponin T with distinct striated fibers (Figure 2). Over the course of 5 DIV, the cardiac myocyte aggregates spread out and organized into a 3D bell shaped form with a height of around 90μm (Figure 2, D-H).

**Figure 2.**
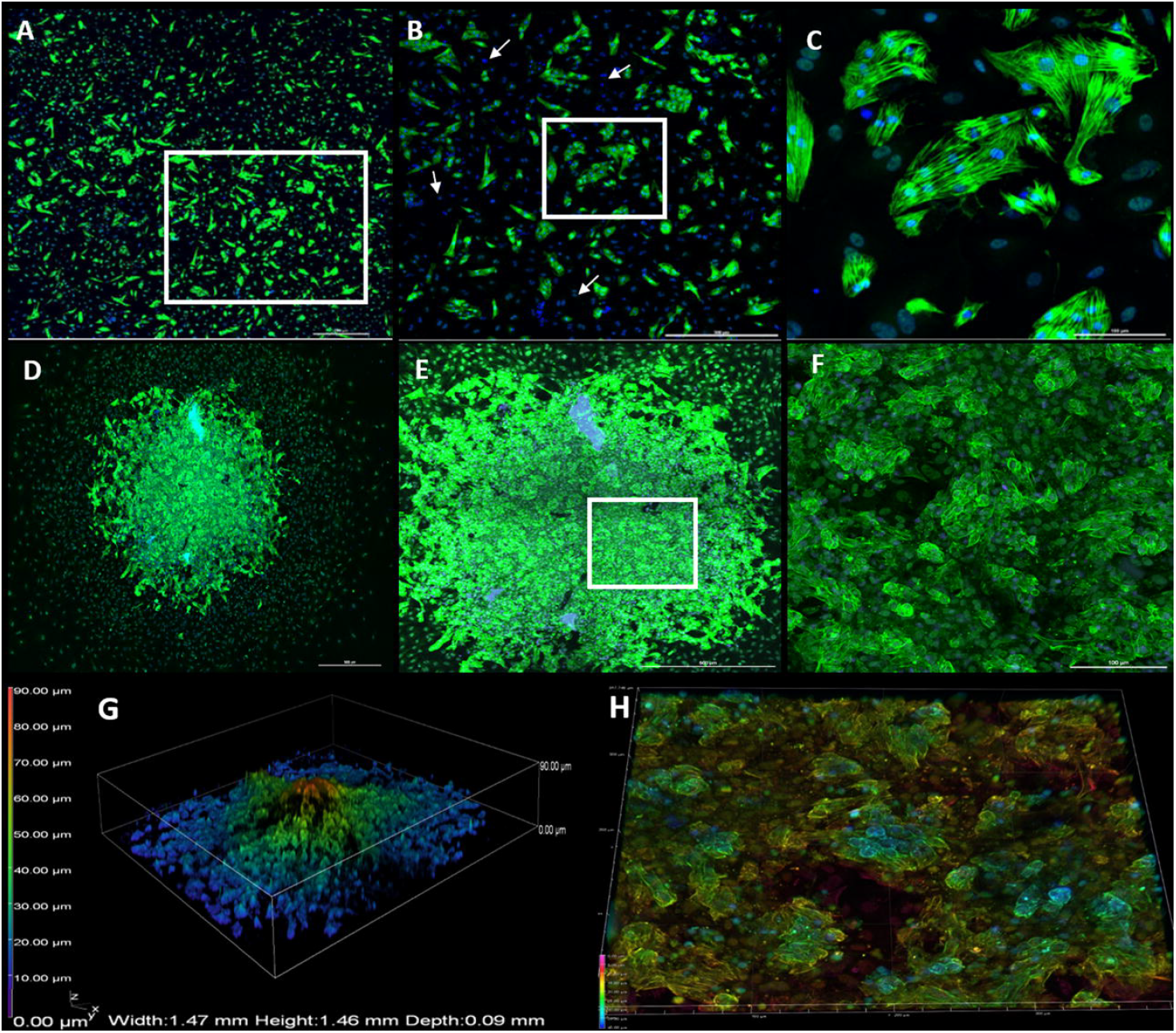
Cell morphology in dissociated and aggregated cultures of cardiac myocytes. Dissociated cultures of cardiac cells expressed cardiac troponin-T (green) (A-C) and striated fibers were observed in individual cells (C). Cell nuclei were stained with Hoechst (blue). Cells staining positive for Hoechst and negative for cardiac troponin-T were of non-cardiac origin (some are indicated by arrows). Cardiac aggregates comprised of a cluster of myocytes in the core region leading to strong signal for cardiac troponin-T and relatively separate cell population around the periphery resulting in milder fluorescence (D). A closer look at the core of the aggregate reveals how myocytes are closely interconnected and growing on top of one another leading to the formation of the 3D aggregate structure (E-F). Volumetric view of the same aggregate in depth coded scale depicting a bell-shaped distribution of cells where blue is closer to the plane and red is at a height of around 90μm (G). Volumetric view of the aggregate core region showing overlay of cells leading to a height of around 40 μm (blue regions) whereas more planar regions are depicted by green (20-25 μm) (H).

Both dissociated and aggregated cardiac myocytes were successfully cultured on MEAs (Figure 3). For the myocyte aggregate cultures, four individual aggregates were plated on each MEA and by 5 DIV these tended to fuse into one larger aggregate. As expected, both dissociated and aggregated cardiac myocytes generated spontaneous contractions that were recorded by the contacts of the MEA (Figure 4A and 4C) however only aggregated cultures generated field potentials discernable at a distance (Figure 4B and 4D). Contractions were both manifest as regular bursting activity recorded from the multi-electrode arrays upon which they were grown, and were visible under light microscopy (Supplementary Movies 1 and 2).

**Figure 3.**
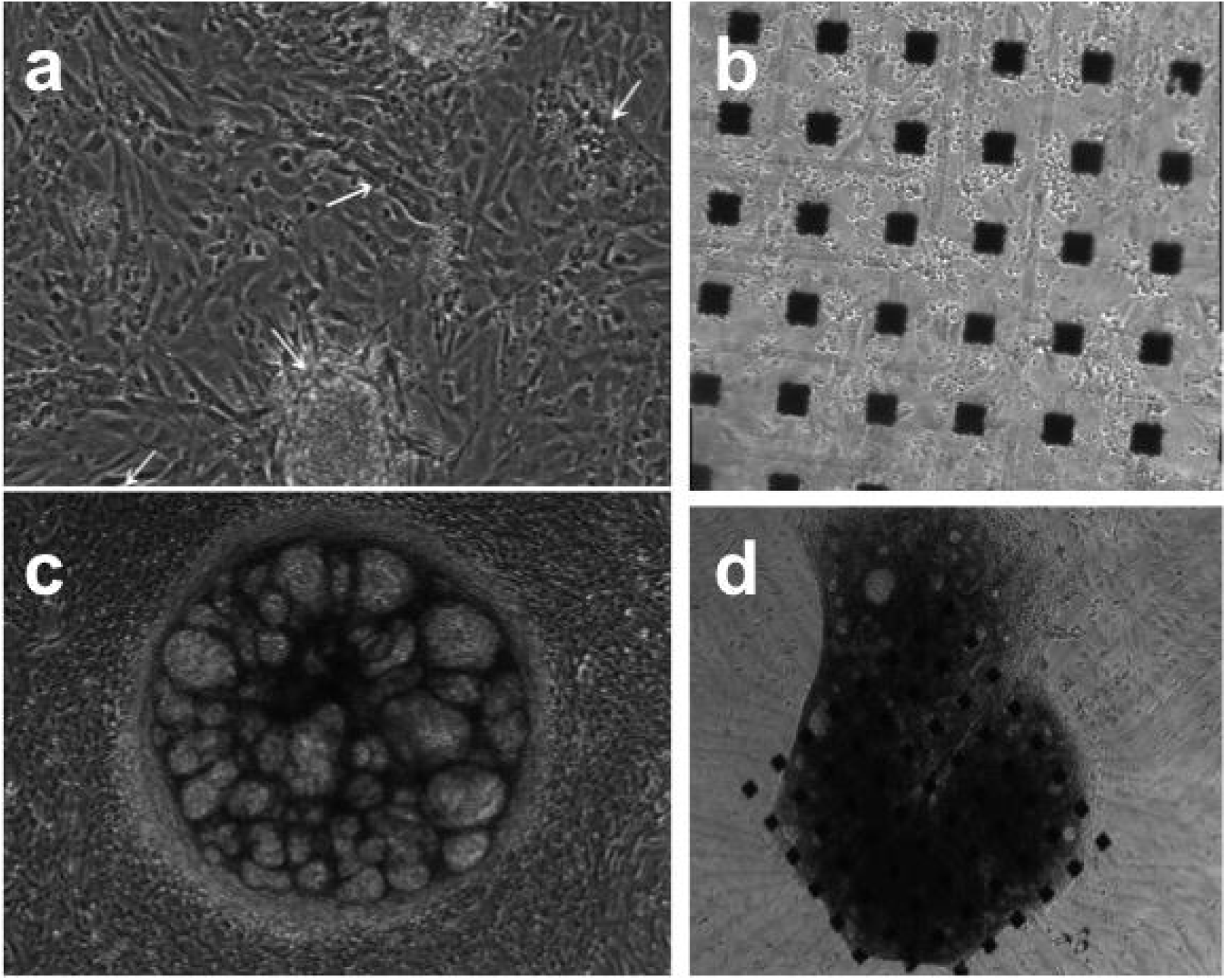
Cardiac myocyte morphology grown dissociated versus aggregated in culture. Dissociated cardiac myocytes grown on polystyrene (a) and atop an MEA (b). Cardiac myocyte aggregates on polystyrene (c) and atop an MEA (d).

**Figure 4.**
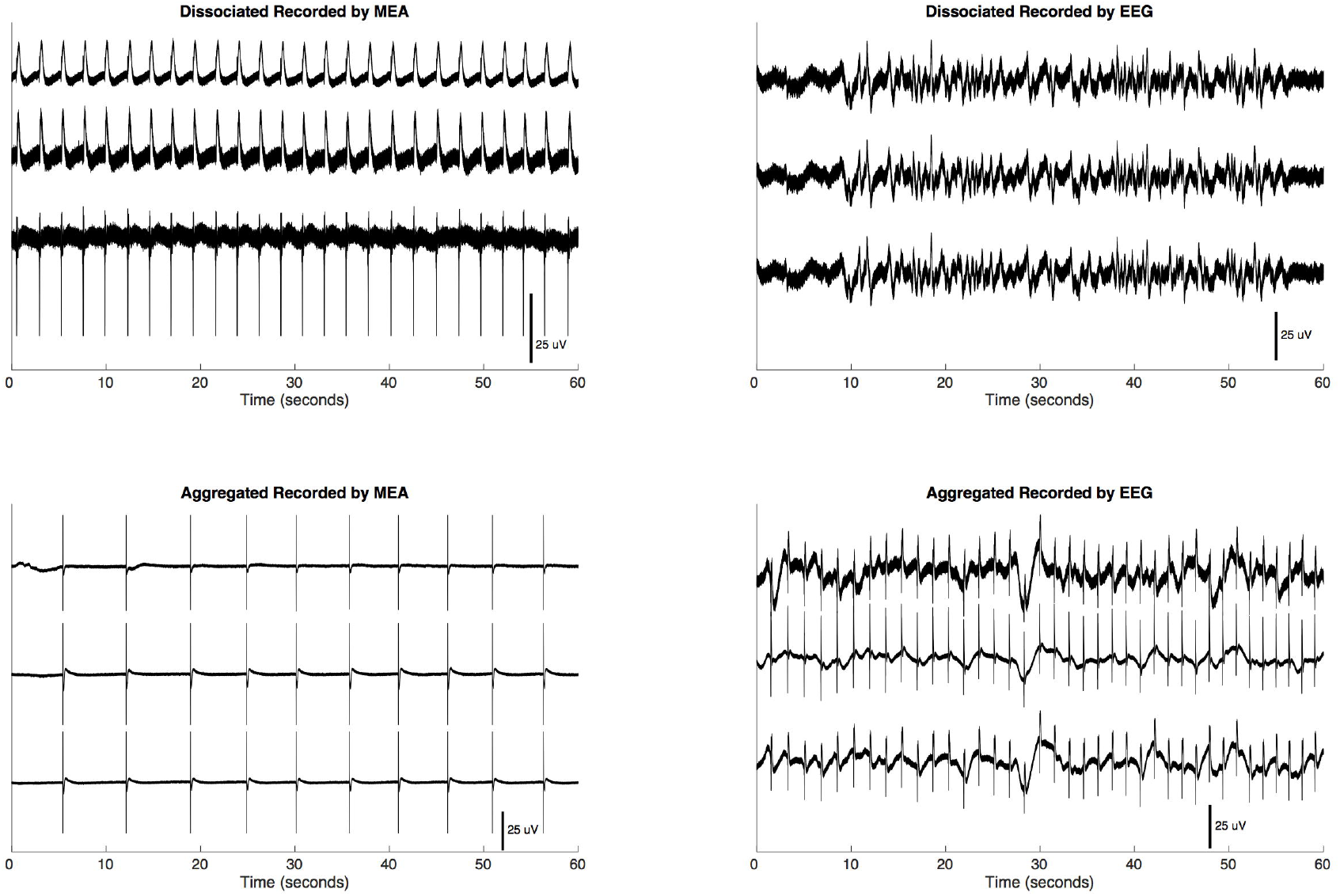
Spontaneous contraction potentials recorded by multi-electrode arrays and mouse EEG of dissociated and aggregated cardiac myocyte cultures. Dissociated cardiac myocytes spontaneously generate contraction potentials as recorded from the underlying MEA (A). Even when beating were confirmed by video microscopy (see Supplementary Movie 1), no commensurate local field reflecting the periodic contraction was observed at a distance by the mouse EEG (B). The MEA recorded well-defined contraction potentials (C) generated by aggregated cardiac myocytes. Even at a distance of several millimeters, periodic contraction potentials could be discerned from the mouse EEG floating above aggregated cardiac myocytes (D).

Distant channel recordings (using the floating mouse EEG) either mirrored or showed the low frequency components of cardiac myocyte aggregate potentials. These recordings were more consistent across distant channels compared to recordings by electrodes in contact (MEA) (Figure 5). To examine how spectral power varied across channels across time, continuously sampled data was convolved with Morlet wavelets with a width of 6 [11], in 1 Hz bands ranging from 1 to 30 Hz, and a subset of bands are shown in (Figure 6). This power spectral analysis of signals recorded simultaneously from the MEA and the mouse EEG revealed several findings: first, low frequency power was of a greater magnitude in the mouse EEG than in the MEA; second, low frequency power fluctuations were uniform across the mouse EEG channels and were heterogeneous across the MEA channels; and third, at higher frequencies, the power fluctuations became more uniform across the MEA channels and hence more similar to those on the distance mouse EEG channels.

**Figure 5.**
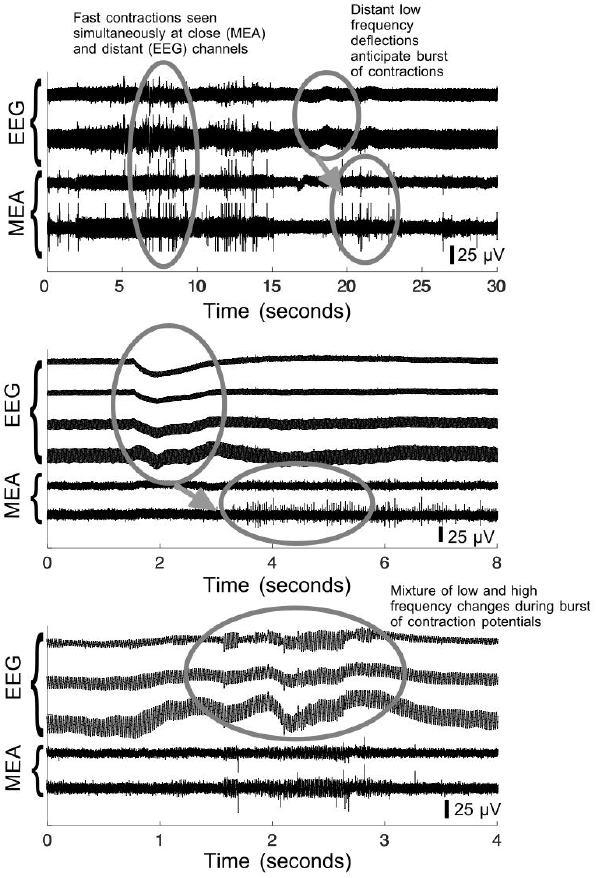
Simultaneous recordings of cardiac myocyte aggregates by underlying substrate MEA and floating EEG. Discrete myocyte potentials are seen on the MEA channels and some of these potentials appear to be broadcast to the EEG (seconds 5 to 15 of top panel), while in other cases, the aggregate appears to function as a low-pass filter with low frequency fluctuations in the remote EEG signal anticipating spike-like contractions recorded in the MEA (seconds 17 to 23 of top panel). This phenomenon of a low frequency potential anticipating a burst of contraction activity is seen again in the middle panel. At other times low and high frequency activity combines in a complex manner at the EEG electrodes relative to the activity recorded at the MEA (bottom panel).

**Figure 6.**
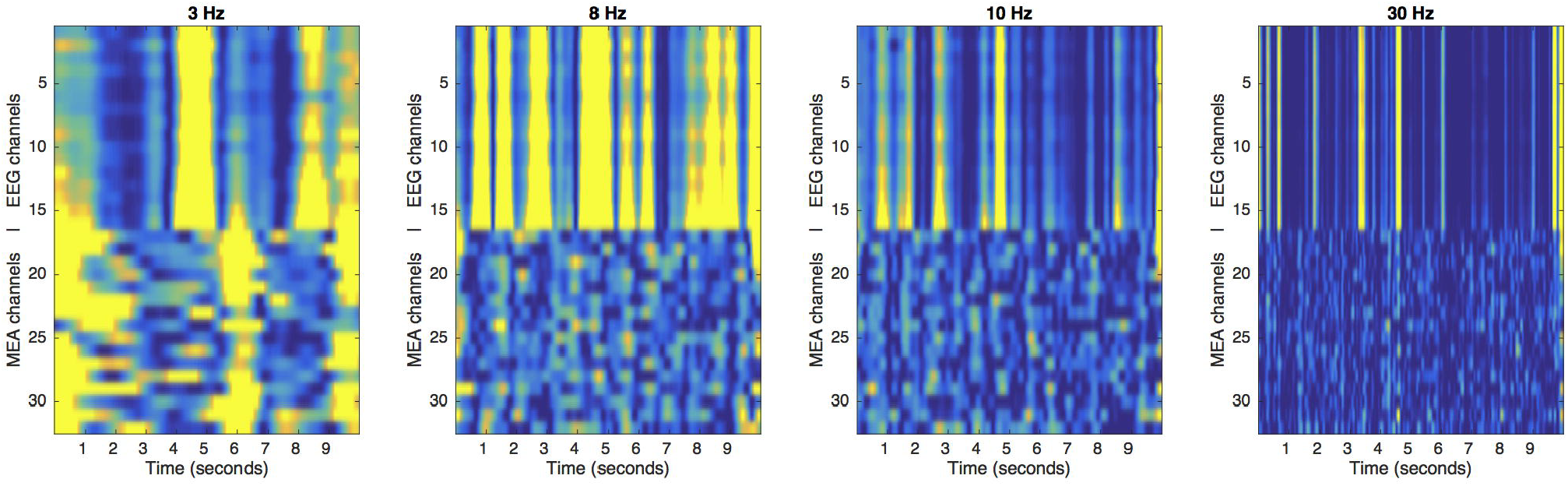
Cardiac myocyte aggregate potential spectral power simultaneously recorded by MEA and floating mouse EEG electrodes. Spectral power was computed with wavelet convolution for four arbitrarily selected distinct 1-Hz wide frequency bands (3, 8, 10, 30 Hz), across 16 channels each of electrodes in contact with the cardiac myocyte aggregates (MEA) and of 16 electrodes 3 millimeters away in culture media (EEG). Overall, spectral power was higher and more uniform across EEG channels than at the MEA. The images share a common color map scheme where dark blue represents 2x10^4^ and yellow represents 2x10^14^ in decibels of power.

By periodically siphoning off small volumes of media (0.4 to 1.2 mL), we were able to decrease the distance from the floating mouse EEG from the aggregated cardiac myocytes (from 0.4 to 1.2 mm).

A fourth-order Butterworth bandpass filter (3 to 2000 Hz) was applied. The amplitude of the signals increased accordingly (Figure 7, left panel). Voltage appeared to follow an exponential decay relative to distance (Figure 7, right panel).

**Figure 7.**
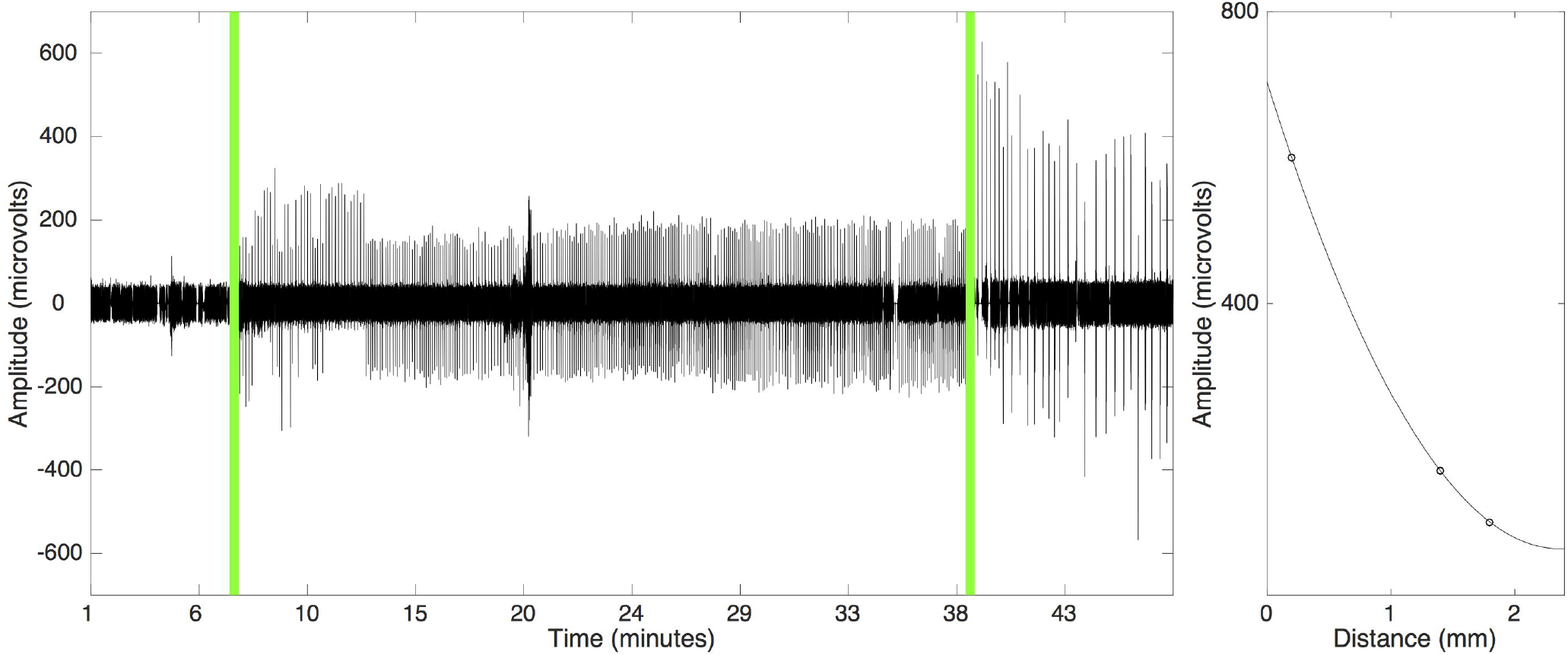
Myocyte aggregate potential amplitude increases by bringing floating electrodes into closer approximation. The left panel shows a recording made using one mouse EEG channel floating above a cardiac myocyte aggregate is shown: a fourth-order Butterworth band pass filter (3 to 2000 Hz) was applied. By removing discrete volumes of media with a micropipette (0.4 to 1.2 mL), the floating mouse EEG could be brought into closer physical proximity (0.4 to 1.2 mm) to the cardiac myocyte aggregates. This procedure increased the amplitude of the potentials recorded, and this effect was most notable in signals place through a bandpass filter. The right-hand panel shows peak contraction voltage recorded by the floating EEG channel as a function of distance from the aggregate; a second-order polynomial is superimposed to illustrate the exponential decay.

The contraction rate of the cardiac myocytes increased roughly 4-fold (from 10-14/min to 40-50/min) with the addition of isoproterenol hydrochloride (noradrenaline) to the media (Figure 8). While the activity of the aggregate initially slowed with each addition of fluid, it quickly recovered spontaneous contractions and additional micro-volumes appeared to help circulate the isoproterenol to accelerate contraction rate.

**Figure 8.**
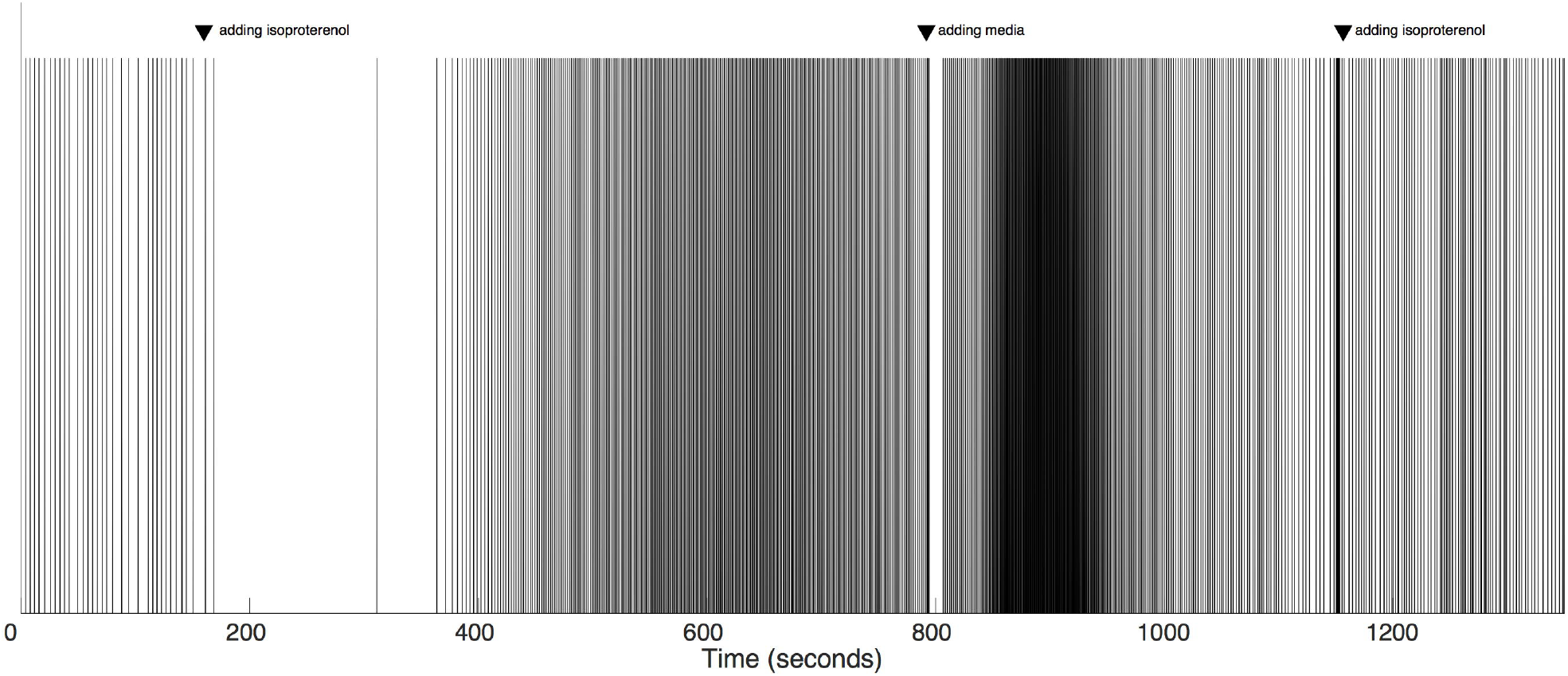
Isoproterenol (noradrenaline) increases the spontaneous contraction rate of the aggregated cardiac myocytes. A recording made at one mouse EEG channel floating above a cardiac myocyte aggregate is shown. The first addition of isoproterenol initially caused quiescence followed by spontaneous activity at a higher contraction rate. When rate started to slow, additional media was added to promote circulation of the isoproterenol over the aggregate. When rate began to decline again, isoproterenol was added again. A fourth-order Butterworth band pass filter (3 to 2000 Hz) was applied. To highlight contraction rate, only signals three standard deviations above the mean are shown. To compensate for voltage amplitude changes with the addition of media (see Figure 7), data was re-normalized for each time window following the addition of any isoproterenol or media.

## 4 Discussion

The pyramidal well-based “forced aggregation” technique was effective at consistently generating 3D aggregates of cardiac myocytes with a diameter ranging from 500 – 800 μm (with 500 μm corresponding to ~0.065 μm^3^). An analogous approach has been used effectively for cardiac differentiation in human pluripotent stem cells [12]. We demonstrated that cardiac myocytes, when grown in forced aggregates, spontaneously generated regular contractions, and that these myopotentials can be detected several millimeters away from electrodes floating in the media above the cells. Interestingly, when grown as 3D aggregates, the cardiac field potentials were robust and did not require averaging to enhance signal quality [13]. In cardiac aggregates individual myocytes were observed to be growing on top of each other and each cell connected through gap junctions form a cardiac syncytium (Figure 2, G-H). This resulted in the entire aggregate beating as a single unit which we believe might contribute to the ability of such engineered aggregates to broadcast signals. In planar culture of dissociated cardiac myocytes, while electrical activity was discernable on MEAs in direct contact with the cells, we were not able to record field potentials at a distance.

We found that the aggregates consisted of mainly cardiac myocytes expressing cardiac troponin T in the core and around the periphery. However, dissociated cells had a significant presence of cells negative for cardiac troponin-T (Figure 2, A-C) which we believe could be fibroblasts resulting from primary culture of embryonic hearts. Being “non-excitable” cells, the fibroblasts have a direct impact on the electrical properties of cardiac myocytes [14]. The presence of fibroblasts among cardiac myocytes can act as an insulated barrier or “current sink” preventing electrical coupling between individual myocytes as evidenced from local anisotropy and cardiac arrhythmia resulting from interstitial fibrosis caused during myocardial ischemia and hypertrophy [14]–[16].

Interestingly, 3D culture of cardiac myocytes at high density appeared to attenuate growth of non-myocyte cells (e.g. presumably fibroblasts) and facilitated myocyte maturation [17], [18]. Our method of forced aggregation allows the seeding of around 10^5^ cells into one aggregate (500-800μm in diameter) thereby resulting in a very high density of myocytes when compared to 2D planar cultures. We believe this is the principal reason why our 2D cultures have significantly more fibroblasts than aggregates even when the source population is the same in both cases. Minimal presence of troponin-negative putative fibroblasts in cardiac aggregates could be an additional contributor to synchronous beating and generation of strong local field potentials that can propagate up to several millimeters away.

Just as muscle tissue “broadcasts” compound action potentials, whether the cardiac electrocardiogram or skeletal electromyograms, so too did these micro-scale 3D aggregates of cardiac myocytes generate “micro” compound-action-potentials detectable at a distance. As expected, the amplitude of these local field potentials varied with distance. We speculate that the heterogeneous pattern of low frequency power fluctuations across the electrodes immediately in contact with the myocytes (on the surface MEA) reflects multiple, smaller local fields while the homogenous, and higher amplitude, pattern of low frequency power fluctuations across the distant electrodes imply that the aggregates volumetrically summate low frequency power. From a neuroprosthetic decoding perspective, the fact that the higher frequency power fluctuations recorded at a distance matched those recorded in contact with the myocytes is reassuring since intended movement signals can be decoded with greater accuracy from higher frequency local field potential signals [19][20].

Just as “living electrodes” (miniature hydrogel micro-columns seeded with autologous, genetically modified neurons) have been engineered to overcome the biocompatibility, biostability and reliability limitations of rigid silicon probes implanted into parenchyma [9], [21]–[23], we propose myocyte aggregates could form a component of “living amplifiers” in which the neural activity could be recorded, amplified and broadcast out of the brain using entirely biological components [1], [3], [24]. Even if synthetic components were ultimately required, they could be deployed in the intraosseous or subgaleal spaces, or completely externally on the skin. By not having foreign bodies in constant breach of the dura and skull, this approach could improve safety by reducing infection risk, and enhance efficacy by providing a more reliable recording system. Just as physician-engineer teams have deployed skeletal muscle tissue to act as a living biopotential amplifier to facilitate neuroprosthetic decoding, so too could this approach be extended to a microscopic scale for direct cerebral recording. To achieve this feat, additional issues must be addressed. First, a reliable way to connect the brain to myocyte “living amplifiers” must be identified. Surprisingly, both Schwann cells and skeletal myocytes can survive chronically when transplanted into the mammalian cerebral cortex [25], [26].

Those earlier investigations were exploring the putative trophic role of these peripheral cells, and it remains unknown if host parenchymal neurons would ever spontaneously generate functional neuromuscular junctions onto such myocytes and if Schwann or other support cells would promote this connection. A “living electrode” [27] or a cerebral organoid [28]–[30] could enable host neurons to interconnect to neurons within these engineered constructs and myocyte aggregates could in turn be grown in or atop these constructs (Figure 9). Since the goal of such amplification is to relay neural signals, skeletal myocytes, which are expected to only depolarize upon receiving acetylcholine from presynaptic neurons, may be preferable over the spontaneously active cardiac myocytes. Rather than coaxing cortical neurons with the host parenchyma to form junctions directly onto skeletal myocytes, the “living electrodes” or organoids could be seeded with motor neuron aggregates. If placed within primary motor cortex, motor neurons also provide an additional advantage: since 5% of the primate corticospinal tract synapses directly on to motor neurons in the spinal cord, there may be a latent fetal phenotypic program available that would predispose host layer V Betz cells to sprout axon collaterals to synapse on to the motor neurons in the constructs.

**Figure 9.**
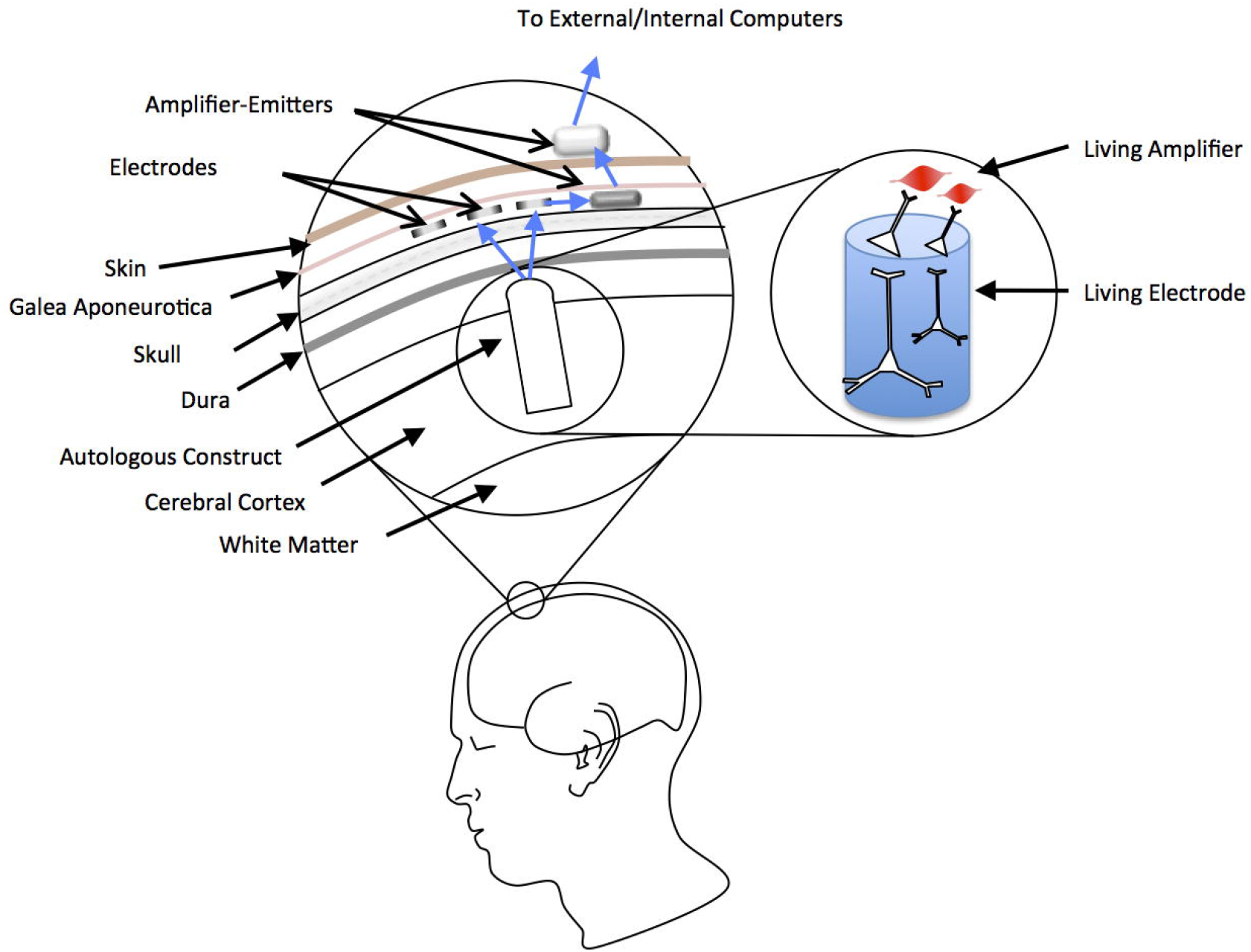
A hypothetical “living electrode”-“living amplifier” scenario for neuromotor decoding in a next-generation brain-computer interface neuromotor decoding. An engineered construct, seeded with autologous induced pluripotent stem cells transformed into neurons and myocytes, would be implanted into the cerebral cortex. The myocyte (or electrocyte) aggregate, would be innervated by neurons ultimately driven by host neuron firing rates, and would generate local field potentials that could be recorded through the dura. Implanted electrodes (such as in strips or grids) implanted in the skull itself, in the subgaleal space, subdermally, or even externally on the skin surface, would record these myopotentials and in turn relay them to an attached emitter that would in turn broadcast them wirelessly either to a subsequent emitter.

In addition to amplifying neural signals, myocytes also function as leaky integrators and low-pass filters. These properties offer an additional crucial engineering advantage: rather than having to record single neuron activity at 30 kHz to decipher and classify action potential waveforms, the myo-amplification process inherently “sorts and counts” spikes, such that subsequent synthetic recording devices only need to sample at 2000 Hz. When faced with numerous channels in parallel, this lower sampling rate affords considerable power and processing advantages, and indeed devices to sample and wirelessly transmit multiple channels of data at this sampling rate already are available for human clinical use [31], [32].

While deploying biopsied skeletal muscle to cap avulsed ends of brachial plexus branches may be feasible in the residual limb of an amputee, there may not be enough physical space or flexibility on the cortical surface to allow multiple contractile elements in a micro-scale “living amplifier” system. Although mammalian myocytes can be rendered non-contractile via addition of a myosin inhibitor (e.g. blebbistatin) *in vitro*, this approach will not function chronically *in vivo*. Several species of fish have evolved tissues and organs, invariably derived from myocyte lineages, to stun prey and communicate with conspecifics [33], [34]. Just as evolution has selected for these so-called electrocytes in fish to build and deliver electrical current at macroscopic scales through water, discarding contractility in the process, in future studies we can ask whether we could artificially convert a mammalian myocyte an electrocyte. It remains unknown if, via the application of small interfering ribonucleic acids, growth factors and CRISPR, whether such a conversion were possible. The ability of electric fish to generate, store and discharge current arises not merely from the presence of electrocytes, but from their stacked arrangement in series at a tissue and organ scale [35]. Hence we may need to purposefully recapitulate these architectural elements, such as arrangement of electroyctes into columns of small disks, in a putative “living amplifier” to enable reliable broadcast of signals through the dura and skull. In our current work, the engineered cardiac aggregates have high cell density stacked within a 3D spherical space that could be mimicking the stacking of electrocytes. Because electric field magnitude decreases in accordance with the inverse square law, one would not expect “living amplifiers” could be designed to emit signals through the air; the purpose instead would be to reach synthetic electrodes that were linked to radio emitters that would in turn broadcast signals to computers or device effectors elsewhere in the body (e.g., epidural spinal stimulators, implantable pumps, robotic prosthetic limbs). As electrocytes were evolved for fish predation and communication in water, it is unclear if animal cells could ever be coaxed also into “living radio emitters” capable of generating kilohertz millielectronvolt signals. Thankfully, low-cost, high-channel radio emitters already exist, both that can be worn non-invasively, and implanted, such as used in cochlear and epiretinal implants [31], [32], [36], [37] (although in this case, signals would be broadcasted out rather than relaying stimulation instructions in).

In summary, here we established that approximately 10^5^ cardiac myocytes, grown as a forced aggregate comprising a volume of approximately 0.065 mm^3^, was able to generate microvolt scale local field potentials detectable several millimeters away with medical-grade electrodes. Further work on driving these myocyte aggregates by neural inputs could prepare the foundation for the design of “living amplifier” components of brain-computer interfaces to treat people with paralysis and other types of neurological disease and injury.

## 6 Author Contributions

MDS, SD, DKC: design of experiments; MDS, SD: performance of experiments; KK, LS: culture of cardiac tissue; SD, JB: histology and microscopy; MDS, SD: data analysis, preparation of figures; MDS, SD, DKC: manuscript preparation.

## 8 Acknowledgments

The authors thank Kate Panzer for technical assistance.

